# Uncovering Sleep’s Hippocampal-Cortical Dialogue: The Role of Deltas’ and Spindles’ Cross-Area Synchronization and Ripple Subtypes

**DOI:** 10.1101/2025.03.19.644132

**Authors:** Adrian Aleman-Zapata, Pelin Özsezer, Kopal Agarwal, Irene Navarro-Lobato, Lisa Genzel

## Abstract

Hippocampal ripples, critical for sleep-related memory consolidation, are heterogeneous events with various sources and functions. Here we applied principal component analysis to identify ripple sub-types and relate them to hippocampal-cortical interactions as well as their role in consolidating simple and complex semantic-like memories in rats. Three main ripple types were discovered: *baseline*, l*arge-input*, and *small-input* ripples. *Small-input* ripples, were associated with increased prefrontal cortex to hippocampus connectivity, followed hippocampal delta waves, and were sufficient for simple learning. In contrast, *large-input* ripples exhibited increased hippocampus to prefrontal cortex connectivity, occurred during hippocampal spindles together as a doublet with a *small-input* ripple, and were critical for complex memory consolidation. Finally, learning induced heightened coupling between hippocampal delta and spindle oscillations and their cortical counterparts, consequently leading to an increased synchronization of ripples with cortical oscillations.

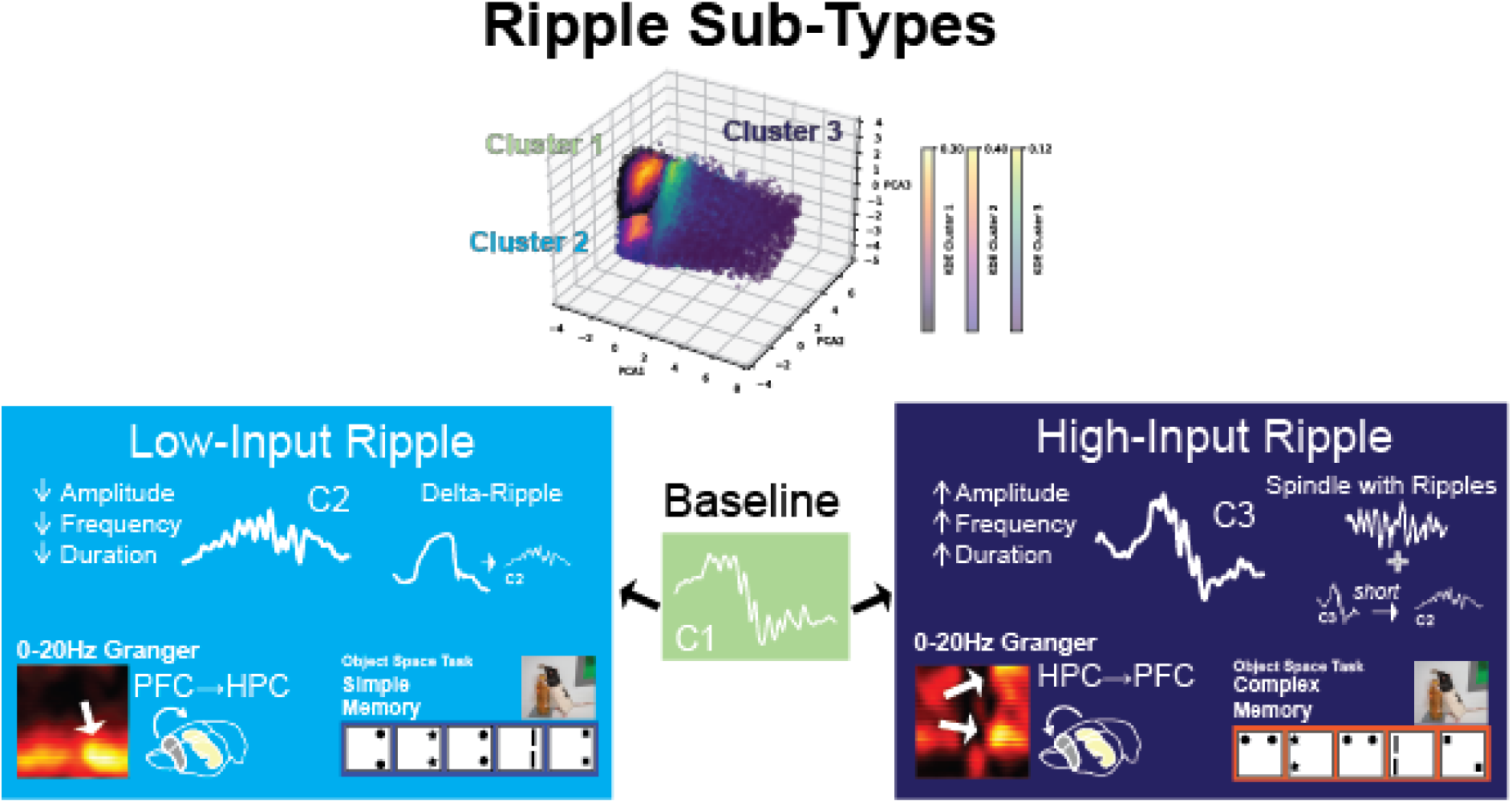

**Significance Statement:** This pioneering, data-driven approach is the first to connect distinct ripple sub-types to precise cross-brain computational states, revealing their role in consolidating various types of memories.

## Introduction

Hippocampal ripple oscillations are critical for memory consolidation [1, 2] and underlie the beneficial effect of sleep on memory [3]. The spiking content in the CA1 hippocampal layer during ripples is influenced by various brain regions [4–6] with inputs from the hippocampal CA3 and CA2 as well as medial entorhinal cortex shown to influence ripple activity [7–9]. However, it remains unknown how ripple types, differing in their primary input sources, correspond to the diverse cognitive processes linked to ripple activity.

Ripples exhibit significant diversity in their characteristics, suggesting a heterogeneous event arising from multiple contributions. Recent studies have demonstrated that unsupervised learning techniques and threshold-based approaches can be employed to disentangle this inherent diversity [10–12]. Using the local field potential trace of the ripple, these methods can differentiate ripple types based on their amplitude, frequency, duration and other features and it has been shown that these ripple types relate to differences in input sources [10, 12].

Previous efforts have successfully identified different types of ripples but have not established their functional relevance. To elucidate this relationship, ripple clustering must be performed in experiments involving significant behavioural manipulations and corresponding readouts. In this study, we detected and clustered ripples occurring during sleep following simple and complex learning conditions, compared to home cage controls, using the Object Space Task [13]. Further, we included a condition manipulating cortical plasticity to elucidate top-down cortical control of ripple types [4].

We identified three types of ripples and investigated their relationship with cortical neuronal firing, oscillation coupling (spindles and delta waves), and contributions to different memory types. Next to the baseline, default ripple, we discovered *small-input* ripples, which were associated with increased prefrontal cortex to hippocampus connectivity, followed hippocampal delta waves, and were sufficient for simple learning. In contrast, *large-input* ripples exhibited increased hippocampus to prefrontal cortex connectivity, occurred during hippocampus spindles, and were critical for complex memory consolidation. Both types of ripples occurred in bursts of multiple ripple events after complex learning, facilitating hippocampal-cortical dialogue.

## Results

### Identifying Ripple Subtypes with Principle Component Analysis

To uncover different ripple subtypes, various ripple features were computed per ripple from its raw trace (Fig. 1A), and a principal component analysis was applied to the extracted features (PCA, Fig. 1B, C). A Gaussian mixture model (GMM) clustering algorithm was applied on the three PCA dimensions, which accounted for most of the variance of the data (∼90%). Four different clusters were identified, of which the final cluster contained very few events and was discarded as noise (Fig 1D). Figure 1E shows both individual examples as well as median values for the remaining three clusters. Interestingly, cluster 2 (C2) was distinct from the other two clusters as it lacked a slow frequency deflection in the signal. This deflection in C1 and C3 may reflect the volume-conducted sharp-wave component originating in other hippocampal layers. C2 also differed from the other clusters exhibiting a small rise after the initial deflection in the prelimbic cortex right before the ripple event (Fig. 1E right).

**Fig. 1.**
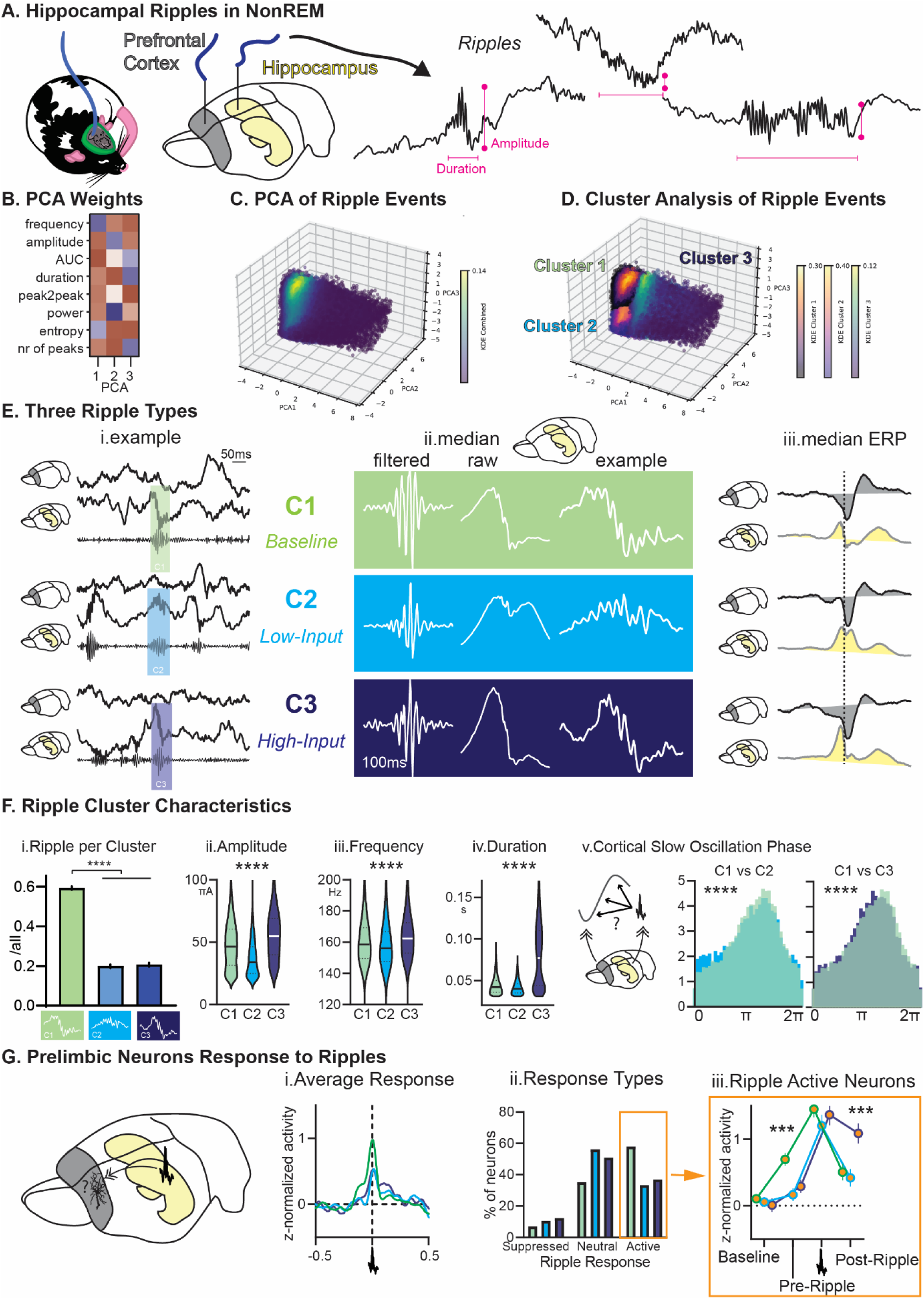
Ripple Subtypes: **A.** We recorded sleep and detected ripples in the hippocampus. Ripples come in many shapes and forms. **B.** features and their weights in each PC **C.** PCA on ripple characteristics showed 3 main clusters (**D.**) determined with a GMM clustering algorithm and evaluated with the Caliksnki-Harabasz index. See SVideo 1 **E.** Median and example traces for each cluster. C1 and C3 are noticeable with larger deflection potentially volume conducted sharp-wave. Traces have been normalized for visualization, showing similar waveforms for C1 and C3. However, non-normalized amplitudes were used in the PCA, and results indicate that C3 ripples had higher amplitudes compared to C1. **F.** Characteristics for ripples from each cluster. Most ripples were part of C1 (i, including PT periods rmANOVA type F_2,186_=404.1 p<0.0001). Subtypes differed in amplitude (ii, including events Kruskal-Wallis=6427 p<0.0001), frequency (iii, including events Kruskal-Wallis=1165 p<0.0001), duration (iv, including events Kruskal-Wallis=11464 p<0.0001), and Slow Oscillations phase coupling (v, each KS- D p<0.0001). cluster 1 green, cluster 2 light blue, cluster 3 dark blue **G.** Response of prelimbic neurons to ripple subtypes. Cluster 1 had a higher ripple response in prelimbic neural activity (rmANOVA cluster*time F_196,10976_=3.1 p<0.001), marginally more ripple-active neurons (Chi-square 8.425 p=0.077) and higher average activity of ripple-active neurons before the ripple (rmANOVA cluster*time F_6,210_=4.7 p<0.001, cluster 1 vs other pre F_1,71_=15.8 p<0.001). Cluster 3 had higher average neuron activity after the ripple (cluster 3 vs other post F_1,69_=14.2 p<0.001). *p<0.05, **p<0.01, ***p<0.001, ****p<0.0001

For each recording period the fraction of ripples corresponding to each cluster was calculated, Cluster 1 was the largest cluster with 60% of ripples in contrast to cluster 2 and 3 with each 20% (Fig. 1F). Interestingly, computing ripple characteristics revealed ripples in cluster 2 to be smaller, slower and shorter than those of the other clusters, and these also were less phase-locked to the cortical slow oscillation. In contrast, cluster 3 ripples were larger, faster, and longer. They were as phased-locked as cluster 1 but occurred earlier in the cortical slow oscillation phase.

Next, we analysed the activity of neurons in the prelimbic cortex around ripples (Fig. 1G). Cluster 1 ripples had a higher average cortical, neuronal response (Fig. 1Gi). More neurons were C1-ripple- responsive (Fig. 1G ii), and these neurons ramped up their activity more before the ripples than seen in other clusters (Fig. 1G iii). In contrast, ripples in C3 came with a higher neuronal response after ripples.

In sum, by applying a PCA to extracted ripple features, three distinct ripple types could be identified. The baseline ripple (here C1) as well as the small-input (here C2) and large-input (here C3) ripple type. C2 and C3 ripples were less common than C1 ripples.

### Hippocampal-Cortical Oscillation Interactions during Ripple Types

Hippocampal ripples are known to interact with cortical regions, but the type of interaction may be specific to ripple types. Thus, we next analysed interactions with slower sleep oscillations i.e. spindles and delta waves.

Ripples are known to be coupled to sleep oscillations such as delta waves and spindles [4, 14–18]. Thus, we detected delta and spindle oscillations in both the prefrontal cortex and hippocampus and identified the fraction of ripples that occurred either in a sequence with a cortical or hippocampal oscillation (delta wave or spindle, before, during, after, Fig. 2A). Ripples in general were more likely to be in a sequence with a hippocampal delta or spindle oscillation than a cortical one (Fig. 2Ai). We split the sequences in types of events, focused on the most common sequences (Fig. 2Aii): Delta followed by a ripple (DR) and ripples occurring during spindles (SwR, Fig.2A ii and iii) [4, 19]. Since cluster 3 had a very large spread of ripple durations (Fig.1 D.iv), we split this cluster into short (<=70ms) and long events (>70ms) for this analysis. Interestingly normalized to rates present in the default C1 ripples, cluster 2 ripples were more likely to occur following a hippocampal delta wave. In contrast, long cluster 3 ripples occurred after hippocampal deltas or during hippocampal spindles, while short cluster 3 ripples were observed during both cortical and hippocampal spindles.

**Fig. 2.**
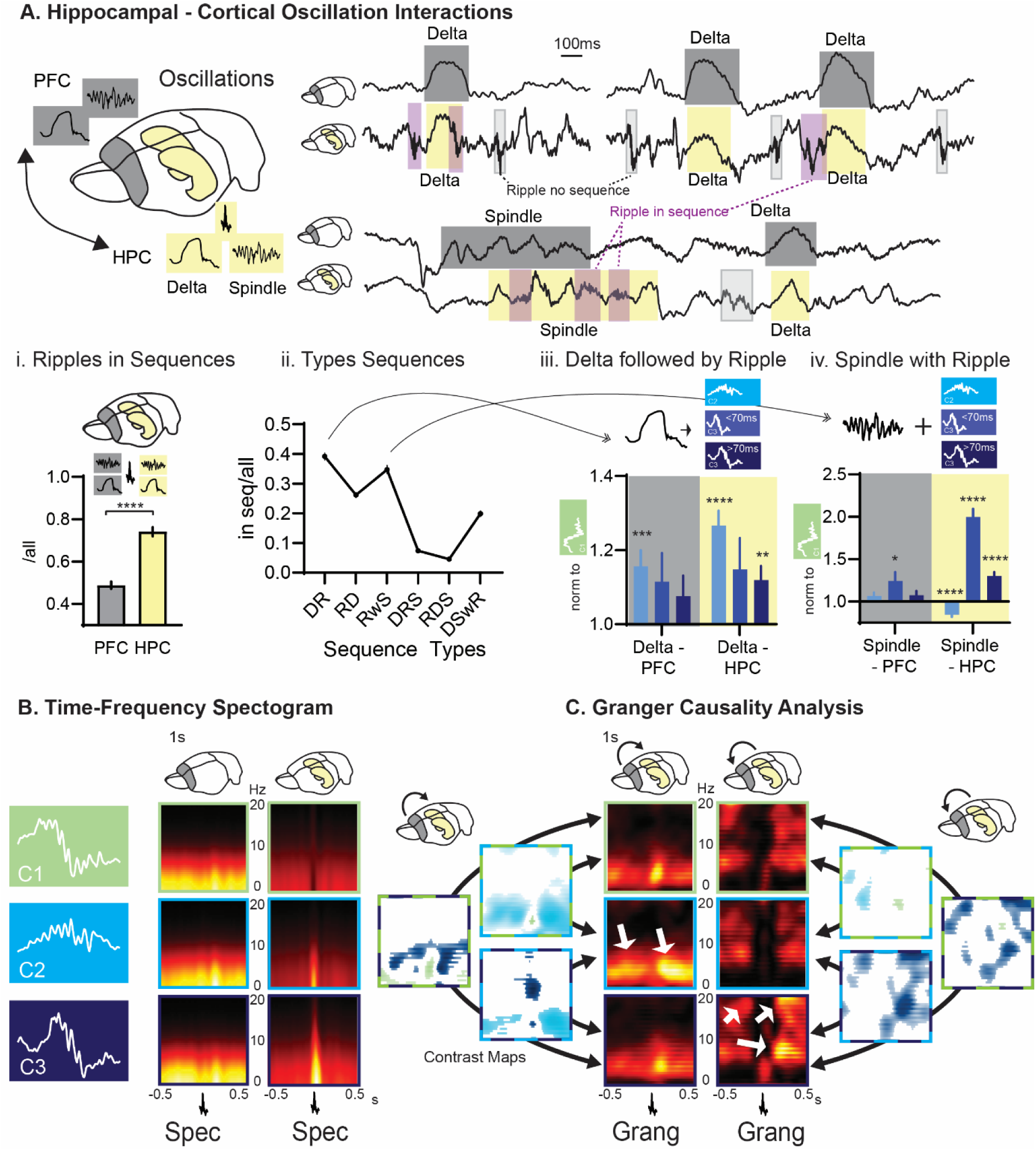
Hippocampal-Cortical Interactions: **B.** Delta and spindle can occur in the prefrontal cortex (PFC) and hippocampus (HPC). **i**. Fraction of ripples occurring in a sequence with a cortical (left) or hippocampal (right) oscillation (t_168_=9.17 p<0.0001). Fraction of ripples occurring either after a delta wave (DR, ii) or during a spindle (SwR, iii) normalized to fraction observed in C1 ripples (one-sample t-test to 1 **p<0.01, ***p<0.001, ****p<0.0001). **B.** Spectrograms (Spec) and **C.** Granger time-frequency analysis (Granger) for prelimbic cortex and hippocampus in lower (0-20Hz, 1s window around ripple, left and right cluster-corrected significant differences between ripple clusters indicated by the arrows) frequency ranges. Clusters differ in the hippocampal sharp-wave (0-4Hz) power, as well as Granger PFC→HPC in the delta and Granger HPC→PFC in the theta (4-8Hz) and spindle/beta (∼20Hz) range (white arrows). Green higher C1, light blue higher C2, dark blue higher C3 for statistical contrast maps. *p<0.05, **p<0.01, ***p<0.001, ****p<0.0001.

Lastly, we computed the spectrograms around the ripples for each cluster and performed time- frequency Granger causality analysis (Fig. 2B and C). We identified significant changes in connectivity with a pixel-based, nonparametric permutation test corrected for multiple comparisons. Cluster 2 was distinguished by higher Granger values PFC→HPC in the upper delta range before as well as after the ripple in comparison to both other clusters. In contrast, C3 ripples showed higher HPC→PFC Granger values before and after the ripple in the theta and spindle/beta range in comparison to both other clusters. C3 ripples also came with higher connectivity for PFC→HPC in 100-300 Hz (Fig. S2). C3 short and long ripples showed the same pattern of connectivity (Fig. S2) In sum, while C2 and C3 ripples resulted in less cortical neuronal firing than C1 ripples, the response was sustained longer after C3 ripples. Interestingly, C2 ripples occur more after hippocampal delta events and short C3 ripples during spindles. Finally, C2 had distinct connectivity for PFC→HPC upper delta range and C3 for HPC→PFC in the theta and spindle range.

### Ripple Multiplets Increase with Complex Learning, Foster a Hippocampal-Cortical Dialogue and Occur during Spindles

After establishing the general properties and cross-brain coupling signatures of each ripple type, we next investigated whether different learning experiences were associated with specific ripple sub- types. Animals were trained in the Object Space Task (Fig. 3A) during which they have the opportunity to explore different object-pairs in an open-field enviroment. As shown previously, the task tests semantic-like memory and animals express the cummulative memory – the statistical distribution of the objects – over trials [4, 13, 19, 20]. We compared a home cage control (remaining in recording box) to different Object Space Task conditions (Fig. 3B): The simple learning condition – same object locations for five trials – and the complex learning conditions – different object locations for five trials. Complex learning consisted of two subconditions: Random (all locations used the same number of times) and Overlapping (one location always has an object, second location changed from trial to trial); however, we have previously shown that these two conditions result in the same consolidation signatures during sleep and present the results combined here [for a more detailed discussion and behavioral results, please see 4]. Interestingly, in both simple and complex learning less C1 ripples were observed, in contrast C3 ripples increased in complex learning (Fig. 3C).

**Fig. 3.**
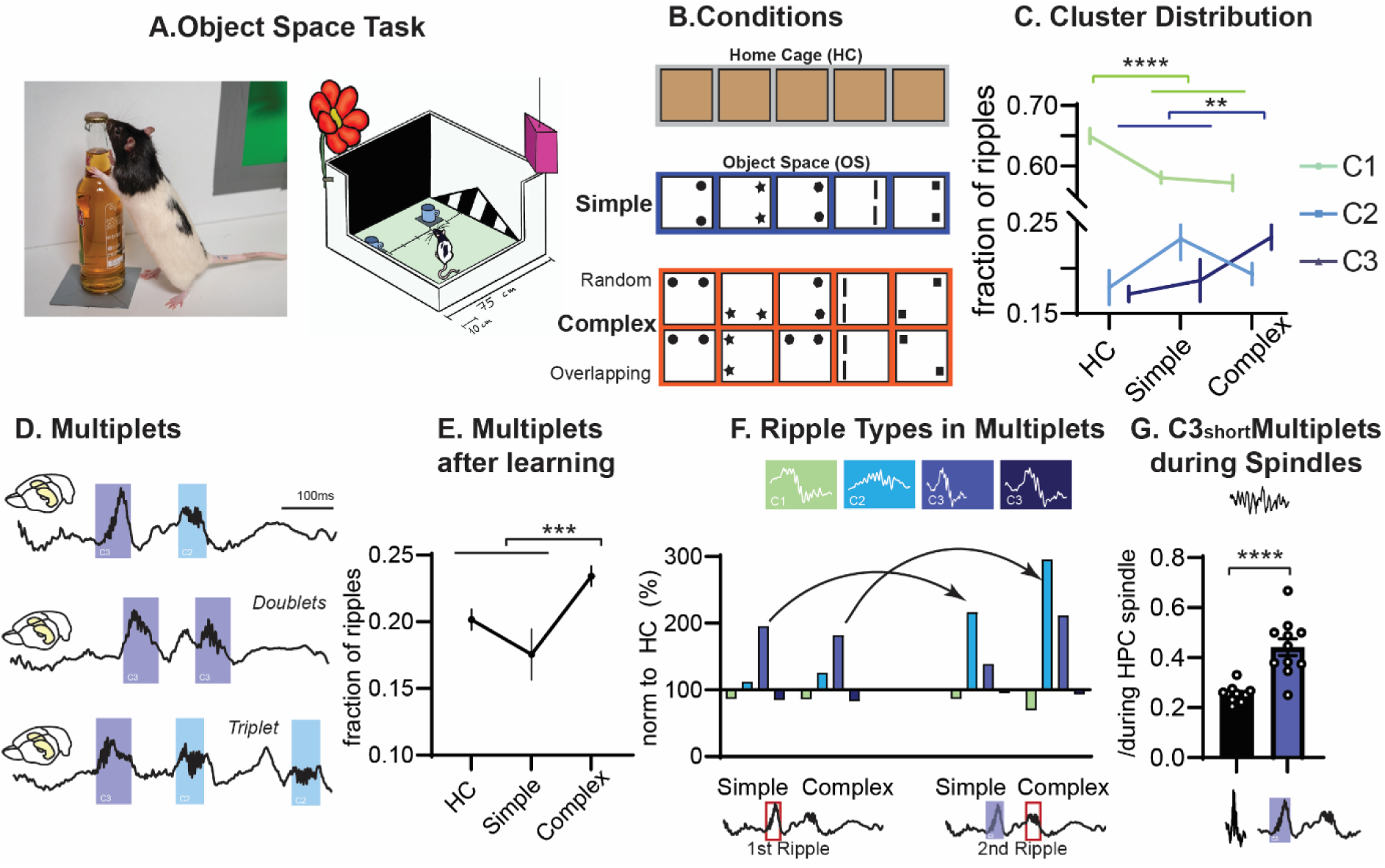
Learning Specific Effects – Multiplets and Spindles. **A.** Object Space Task allows animals to explore objects in an open field. **B** Home Cage (HC) was compared to simple and Complex learning conditions. **C**. Fraction of ripples per PT period classified as C1, 2, or 3 split for different behavioral conditions (2-way ANOVA condition F_2, 246_=0 p>0.99, cluster type F_2, 246_=625.5 p<0.0001, interaction F_4, 246_=7.3 p<0.0001) **D**. Example of ripple multiplets, here with C3 and C3 ripples. **E.** Fraction of ripples that were part of mulitplets split for condition (F_2,82_=7.6 p=0.0009). **F**. Ripple cluster type that was the first ripple of the multiplet (left). Right: Taking short C3 ripple events and testing which type is the second ripple. Each normalized to home cage. **G**. Fraction of C3short-Multiplets and all ripples occuring during spindles (t_21_=5.6 p<0.0001) *p<0.05, **p<0.01, ***p<0.001, ****p<0.0001

Ripples are known to occur in bursts, with multiple ripples in close succession, i.e. doublets or multiplets. Thus next, we tested how many ripples occurred in multiplets and which ripple types contributed to multiplets (Fig. 3D). After complex learning the fraction of ripples part of multiplets increases (Fig.3E). Interestingly, in comparison to home cage, learning – both simple and complex – will shift which types of ripples are part of multiplets. After learning, the first ripple is more likely to be a short C3 ripple and these short C3 ripples are then more likely to be followed by a C2 ripple or another short C3 ripple (Fig. 3F, SFig. 1D). Finally, it has been proposed that multi-plets occur during spindles [21], thus we compared the fraction of C3short-led multiplets that occurred during spindles to general ripple-spindle coupling. C3short-led multiplets were more likely to occur during a spindle than other ripples.

In summary, simple learning is associated with a shift characterized by a decrease in C1 ripples and an increase in C2 ripples, whereas complex learning results in a higher occurrence of C3 ripples. Additionally, complex learning induces more ripple multiplets, which alternate between C3 and C2 ripple events, and occur during spindles.

### Delta and Spindle Cross Brain Coupling and Learning

Next, we investigated the delta and spindle cross-brain coupling and how it related to our learning conditions (Fig. 4A). We saw that hippocampal delta and spindle events were more coupled to their cortical counterparts after learning in comparison to home cage (both simple and complex learning, Fig. 4B). Interestingly, this led to more ripples being part of cortical sequences, while there was no such effect for hippocampal sequences (Fig. 4C, SFig. 1B and C).

**Fig. 4.**
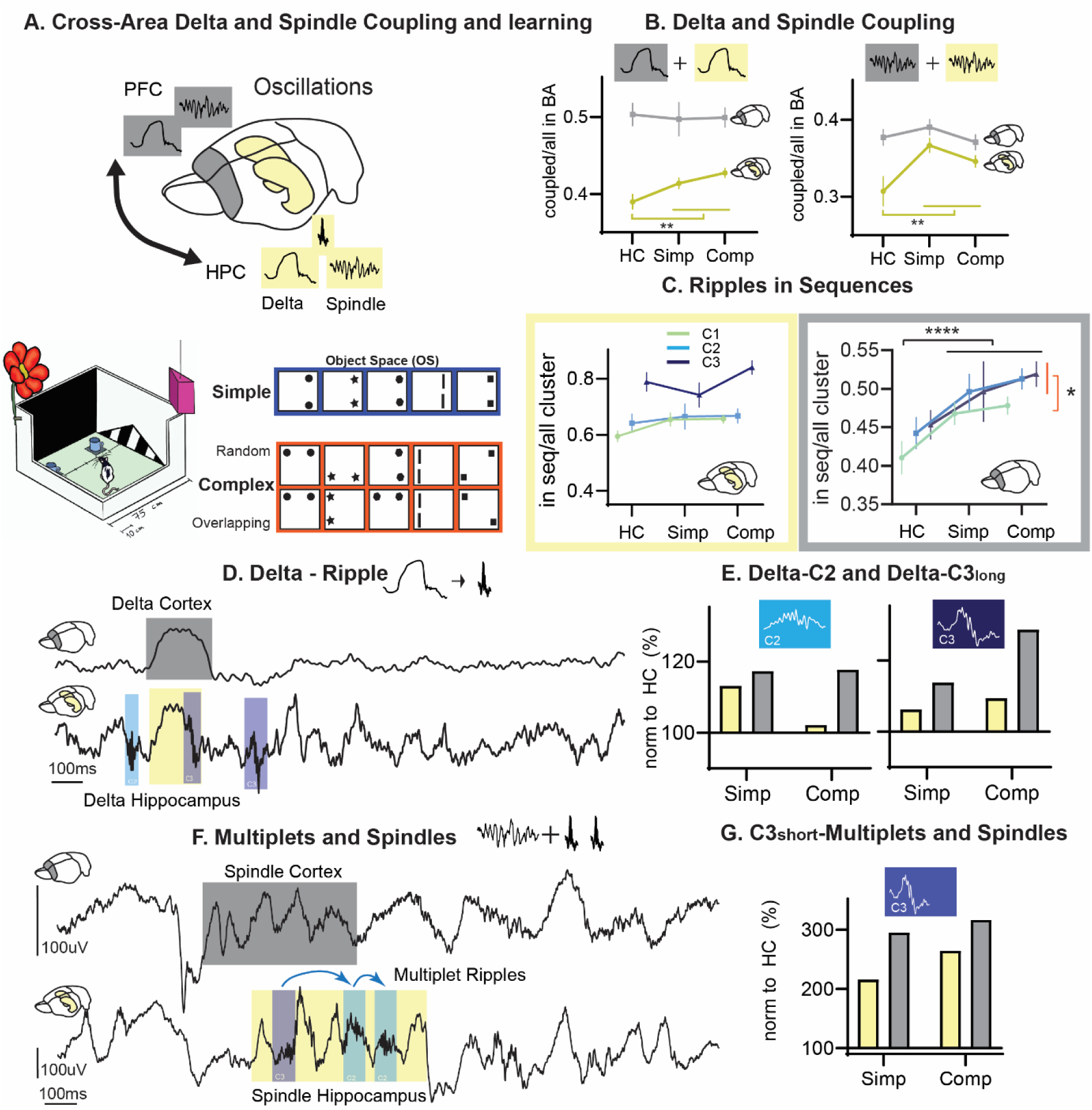
Learning Specific Effects – Sleep Oscillations. **A.** Home Cage (HC) was compared to simple and Complex learning conditions. **B**. Fraction of cortical (grey) and hippocampal (yellow) delta (left, 2-way ANOVA condition F_2, 166_=0.9 p=0.4, brain area F_1, 166_=70.6 p<0.0001, interaction F_4, 166_=1.4 p=0.25) and spindle (right, 2-way ANOVA condition F_2, 166_=4.2 p=0.02, brain area F_1, 166_=17.5 p<0.0001, interaction F_4, 166_=2.3 p=0.1) that occurred coupled to a corresponding event in the other brain area split for behavioral condition **C**. Fraction of ripples occurring in a sequence with a cortical (left, 2-way ANOVA condition F_2, 246_=9.4 p=0.0001, cluster type F_2, 246_=3.0 p=0.0553, interaction F_4, 246_=0.03 p=0.99) or hippocampal (right, 2-way ANOVA condition F_2, 246_=2.4 p=0.09, cluster type F_2, 246_=23.3 p<0.0001, interaction F_4, 246_=1.1 p=0.35) oscillation split for cluster and condition**. D** Example of ripples occurring before and after delta waves **E.** Shows C2 ripples and C3long ripples that occur after delta waves normalized to all ripple count and home cage split for condition (C2 Chi-Square_6_=13.8 p=0.033, C3 long Chi-Square_6_=14.1 p=0.028) **F.** Multiplets occur during spindles **G.** Shows C3short multiplets that occur during a prelimbic (PFC) and hippocampal (HPC) spindle normalized to all ripple count and home cage split for condition (Chi-Square_6_=52.5 p<0.0001) *p<0.05, **p<0.01, ***p<0.001, ****p<0.0001

We demonstrated that C2 and long C3 ripples tended to follow delta waves while C3short multiplets occurred during spindles (Fig. 2B). Next, we investigated if these event sequences were more likely to occur after learning and could show a moderate increase of 20% for the C2 and long C3 ripples following delta waves (Fig. 4 D, E) and a large increase (100-200%) of short C3 ripple multiplets occuring during spindles (Fig. 4 F, G).

In summary, learning enhances cortical-hippocampal coupling (both delta-delta and spindle- spindle), leading to an increase of ripples occurring in cortical sequences.

### Increasing Plasticity in the Prefrontal Cortex Changes Proportion of Ripple Subtypes

To test whether the occurrence of ripple types is under top-down control of the cortex, we artificially increased plasticity in the prelimbic cortex via the overexpression of an established plasticity-enhancer called regulator of G protein signalling 14 of 414 amino acids (RGS14414, Fig. 5A)[22, 23]. The overexpression of RGS14414 is known to lead to increased BNDF and dendritic branching in the targeted area [22, 23] and thereby increase plasticity locally. We have previously shown that this cortical manipulation leads to *overlearning*, where all experiences – including home cage control – induce consolidation signatures and enhanced one-trial learning leads to more interference on semantic-like memory [4]. Using the PCA loadings from the vehicle animals, we could find the same ripple clusters in RGS14 animals (Fig. 5B, SFig 3 and 4). In RGS14 more ripples were part of clusters 2 and 3 (each t).

**Fig. 5.**
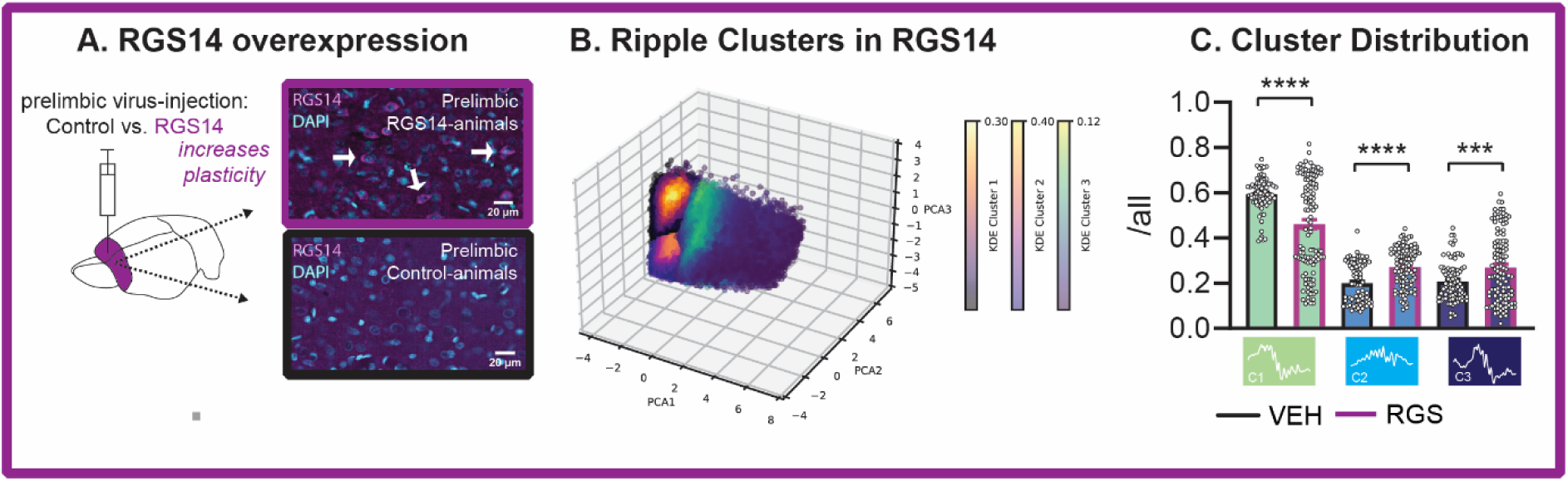
Increasing Cortical Plasticity: **A.** RGS14 was injected into the prelimbic cortex leading to increased plasticity. **B**. In RGS14 animals the same four clusters were found. **C.** Fraction of ripples per cluster (Welsch test each t>3.6) *p<0.05, **p<0.01, ***p<0.001, ****p<0.0001

In sum, increased plasticity in the prelimbic cortex, leads to more C2 and 3 ripples and increased cortical-hippocampal coupling during home cage fitting to the *overlearning* phenomenon [4].

## Discussion

In the present study, we employed an unsupervised learning approach to identify types of ripples characterized by unique features and properties. Consistent with prior research, we distinguished three distinct types of ripples: the baseline ripple (C1, accounting for 60% of ripples) and two variants with *low* and *high-input*, denoted as C2 and C3, respectively (each representing 20% of ripples). A novel aspect of our investigation lies in establishing the functional significance of these ripple subtypes: *low-input* ripples, which follow delta waves and exhibit heightened cortical-hippocampal connectivity, play a role in simple learning. In contrast, *high-input* ripples, which coincide with spindles and demonstrate hippocampal-cortical connectivity, become more prevalent following complex learning. Intriguingly, following complex learning, both ripple types occur alternately in bursts of ripple events, facilitating a dynamic hippocampal-cortical dialogue.

### Comparison to other ripple types

In our dataset, we identified three distinct ripple types. These types exhibited variations in classic features such as amplitude, frequency, and duration. Additionally, they differed in cortical slow oscillation phase-locking and the presence of a potential sharp-wave deflection. These distinctive features enable us to compare our identified ripple types with previously described classifications.

*Low-input* C2 ripples had no potential sharp wave deflection and thus correspond to subtype 4) of Ramirez-Villegas et al. [11], region C in [10] and ripples without sharp-wave in Aleman et al [12]. This cluster of events may not originate from the CA3 region, as the discharge of this area typically generates sharp waves. Instead, it is probable that this type of ripple is generated by other regions such as CA2 or the entorhinal cortex [7, 10–12]. Such ripples were associated with small current source density sinks in the stratum lacunosum moleculare and small sources in the stratum radiatum, importantly the relative contribution of each differed from *baseline* ripples with a relative higher contribution of the lacunosum molculare where the medial entorhinal cortex feeds into CA1 [10]. However, our results provide tantalizing evidence that these ripples are coordinated by the prelimbic cortex – likely mediated via the medial entorhinal cortex through the delta-wave [7] – as seen in the Granger and coupling analysis. These ripples were not strongly associated with the cortical slow oscillation, and the delta waves preceding them were unlikely to be complete downstates, which are the hallmark of true slow oscillations [24]. Therefore, the delta waves observed in this context are likely indicative of local slow wave activity. These intriguing results hint at a mechanism previously ignored. The thalamus would be a candidate to mediate such learning-induced increase in cross-brain coupled local slow wave activity [25]. Interestingly, these types of ripples were more prevalent after simple learning in the Stable condition of the Object Space Task, wherein information regarding object locations is reinforced from one trial to the next. Under such conditions, the cortex can rapidly assimilate the information, potentially exerting an increased influence on ripple activity. We have previously demonstrated that such low-input ripples are sufficient for the consolidation of simple learning in the same task [12, 19].

*High-input* C3 ripples exhibit a more diverse array of features. They stem from a substantial input from CA3, often quantified as sharp-wave sinks in the CA1 stratum radiatum. The magnitude of this input has been demonstrated to correlate with ripple power and frequency [9, 26]. Long ripples were included in this cluster, similar to those reported in wake [27] and sleep [19]. Ramirez-Villegas, Logothetis [11] referred to these ripples in anesthetized rhesus monkeys as “*classical*” ripples (subtypes 2 and 3). In their article, they described them as ripples, which coincide or precede closely the sharp-wave dendritic depolarization peak and have larger sharp-wave amplitudes as well as higher ripple frequencies. These ripples came with larger cortical response in BOLD signal and more subcortical suppression. In mice, Nitzan, Swanson [28] elucidated that ripple power correlated positively with the sharp-wave sink magnitude, ripple duration, ripple frequency and the synchrony of CA1 and CA3 neurons. Our data confirms this with the larger spectral power observed in the sharp-wave frequency band of C3 ripples, which were bigger, faster and longer compared to the other ripple clusters. Moreover, long-duration ripples showed a lower E/I ratio in the hippocampus and higher hippocampal neuronal activity compared to short ripples and random NonREM baselines [19]. These characteristics create better conditions for the ripple to occur and be maintained [9, 29]. These ripples also came with an increase in HPC→PFC connectivity in the theta and spindle/beta range especially after the ripple event itself and occurred during spindles. These two hot-spots in the time-frequency Granger analysis are remarkably similar to cortical spectrograms generated after cue-presentation during sleep in targeted-memory-reactivation (TMR) paradigms [30]. It would be tempting to speculate that both are capturing the same type of event: ripples transmitting information by entrained hippocampal-prelimbic cortex theta and spindle oscillations. Correspondingly, only this ripple type led to extended neuronal firing in the cortex after the ripple.

The final group*, baseline* ripples, had intermediate values in frequency and amplitude compared to *small* and *large-input* ripples. Their durations overlap with the range of durations that *small-input* ripples can have, although the latter subtype tends to be shorter. Baseline ripples are the subtype that occurs the most by default during periods of basal activity and no learning. Hall and Wang [31] proposed that these ripples are generated stochastically through a rigid pathway that forms the backbone of preconfigured information. They hypothesized that this pathway mostly targets the rigid CA1 pyramidal neurons described by Grosmark and Buzsaki [32], which activity didn’t exhibit a change between the pre-learning and post-learning sleep. Moreover, the disruption of this ripple subtype during sleep, unlike other ripple subtypes that are involved in learning simple and complex memories, has been shown to not trigger a homeostatic-like upregulation [33]. This lack of upregulation is expected in baseline ripples given that they are prevalent after routine behaviours, like exploration of familiar environments, that don’t require learning. In our study, less of these ripples were seen after learning. Interestingly, these ripples came with the highest cortical neuronal response. Fittingly, others have also reported that learning comes with sparser coding than default activity [28, 34].

In sum, our ripple variations correspond to known types and reflect different hippocampal-prefrontal computational states. Low- and high-input ripples show the reversed hippocampal-cortical connectivity and respond differently to simple and complex learning inputs.

### Oscillation sequences depending on simple or complex learning

Our ripple types showed shifts in participation in different cortical and hippocampal oscillation sequences (spindles and deltas). Ripples occur before or after a delta-wave [15, 16, 35] as well as before or during spindles [16–18, 36, 37]. We had previously described that when different types of experiences and learning are compared, behaviour-specific oscillation-sequences are observed [3, 4, 19].

We can now add that simple learning in rats also results in more *low-input* C2 ripples, that are smaller, may have a cortical origin and are preceded by delta waves. In contrast, complex learning resulted in more *high-input* C3 ripples, that were diverse in features and could be very long. We previously reported that delta-ripple sequences increased specifically after the complex learning conditions [4]. Later, we replicated this effect and showed that this was especially true for longer ripples [19]. Here we can expand the finding and show, that C3 ripples occur both in delta-ripple sequences as well as during spindles. Specifically, shorter C3 ripples are those that occur during spindles, while long C3 ripples occur after deltas. Previously the delta-ripple sequence was described together with prelimbic reactivations when animals learn complex rules [15]. Our C3 ripples come with bilateral communication with increased HPC→PFC connectivity in theta and spindle/beta and increased PFC→HPC in the fastest frequency ranges. This could be a marker of memory updating, where novel information is transmitted from the hippocampus to the cortex, which would be especially important after complex learning with continuously changing information from trial to trial. The primary effect of both simple and complex learning was an augmentation of hippocampal-cortical coupling of delta and spindle oscillations. While this outcome is not unexpected, our study unveiled a novel finding: ripple types inherently exhibited a default coupling with hippocampal delta and spindle events. Following learning, hippocampal delta and spindle activities became more synchronized with their cortical counterparts. Consequently, hippocampal ripples were more prone to occur in sequences alongside cortical delta and spindle oscillations, facilitating the updating of cortical memory networks.

The anterior thalamic nucleus has been demonstrated to synchronize the cortical slow oscillation and spindle activity [25, 38, 39], suggesting a similar entrainment mechanism for hippocampal oscillations given the known interactions between these brain regions. Consequently, the mechanisms underlying increased hippocampal-cortical interactions after learning likely involve heightened slow frequency coupling mediated by thalamic inputs. These coupled slower frequency events, such as delta and spindles, would facilitate the transmission of information from faster events, such as hippocampal ripples, to a more receptive cortex. The identified ripple types may correspond to different receptive states: C2 ripples indicative of cortical selection of hippocampal output, and C3 ripples representing hippocampal reinforcement of new information.

In sum, we show that learning increases cross-brain coupling of slower frequency events to enable the transmission of information during the ripple.

## Conclusion

Applying a data-driven principal component analysis, we delineated different ripple types: a baseline ripple (C1) and two subtypes (C2 and C3), each characterized by distinct spectral profiles, attributes, and involvement in oscillatory sequences. C2 ripples were indicative of cortical-driven activity, exhibiting an increase following simple learning, while C3 ripples were associated with hippocampal spindle events and showed augmentation after complex learning. Furthermore, learning induced heightened coupling between hippocampal delta and spindle oscillations and their cortical counterparts, consequently leading to an increased synchronization of ripples with cortical events. Thus, our findings demonstrate the utility of data-driven approaches in elucidating the functional roles of ripples in the context of learning.

## Contributions

AAZ analyzed the data and supervised students, PO and KA analyzed the data, INL acquired the data and co-designed the study, LG designed the study and led experiments as well as analysis.

## Acknowledgements

Shekhar Narayanan for extracting spiking activity during ripples. Abdelrahmen Rayan, Anumita Samanta, Jacqueline van der Meij and Federico Stella for advice on the project, the work was supported by an NWO-VIDI grant to LG.

## Methods

### Study design

A total of 8 rats were used in this experiment, which we have previously analyzed with a different emphasise and published as [4]. Animals were first extensively handled for multiple days (at least 3) until they did not flinch when the experimenter touched them (see handling videos on www.genzellab.com). Next, all animals underwent viral-injection surgery (see below), half the animals received RGS14_414_-lentivirus while the other had a vehicle (empty) lentivirus. The 8 electrophysiology animals (4 vehicle, 4 RGS) received a second surgery three weeks after the first one, for hyperdrive implantation. During 2–3-week surgery recovery, tetrodes were slowly lowered to target area before the animals also had habituation and training in the Object Space Task.

### Animals

Three-month-old male Lister Hooded rats weighing between 300-350 g at the experiment start (Charles Rivers, Germany) were used in this study. Rats were pair-housed in conventional eurostandard type IV cages (Techniplastic, UK) in a temperature-controlled (20 + 2 °C) room following a 12 h light/dark cycle with water and food provided *ad libitum*. After lentiviral surgical intervention, animals were single-housed for two days and paired with their cage mates after recovery in rat individually ventilated cages (IVC; Techniplastic, UK). Animals were maintained in their IVC in a barrier room for 14 days before downscaling them to conventional housing conditions. After hyperdrive implantation, rats were single-housed in until the end of the experiment. A total of 8 rats were in the electrophysiological recordings (n=4 per treatment, RGS14_414,_ and vehicle, split one animal per treatment across four cohorts). The behavioral experiments and electrophysiological recordings were performed during the light period (between 9:00-18:00).

All animal procedures were approved by the Central Commissie Dierproeven (CCD) and conducted according to the Experiments on Animals Act (protocol codes, 2016-014-020 and 2016-014- 022).

### RGS14_414_-lentivirus

The RGS14_414_- and vehicle-lentivirus solutions (1.72 x 10^7^ CFU/ml and 2.75 x 10^6^ CFU/ml respectively) were prepared and provided by Dr. Zafaruddin Khan at the University of Malaga (Malaga, Spain)[40] [41]. Briefly, the RGS14_414_ gene (GenBank, AY987041) was cloned into the commercial vector p-LVX DsRed Monomer-C1 (Clontech, France) using DNA recombinant technology. Then, both non-replicant RGS14_414_- and vehicle-lentivirus (empty vector) were prepared and titered using the Lenti-XTM (Clontech, France) according to the manufacturer’s instructions.

The animal’s procedures related to the non-replicant lentiviral solution were approved and carried out in compliance with institutional regulation.

### Tetrode hyperdrive

A customized lightweight tetrode micro-drive was manufactured to implant 10 and 6 movable tetrodes in the prelimbic cortex and hippocampus (HPC), respectively [42–44]. Two separate bundles of #33 polyimide tubes (Professional Plastics,) were prepared: one of 2 columns x 5 rows for prelimbic cortex and 3 X 3 for HPC. The bundles were fixed first to the customized 3D printed cannula and then into the customized 3D printed body drive. The 3D printed cannula was designed according to the Rat Brain Atlas in Stereotaxic Coordinates [45] for the correct placement of the bundles in the areas of interest. Inner tubes (#38 Polyimide tubes; Professional Plastics) were placed inside the outer tubes and glued to the shuttle, which moves through the body spokes thanks to an inox steel screw and a spring CBM011C 08E (Lee spring, Germany). A total of 16 tetrodes were built, twisting four 10 cm polyimide-insulated 12 μm Nickel-Chrome wires (80 turns forward and 40 turns reverse) (Kanthal Precision, Florida) and fused by heat. Tetrodes were loaded in the inner tubes, and their free ends were connected to a customized 64 channels, 24 mm round electrode interface board (EIB) using gold pins (Neuralynx). Previously, 2 NPD dual row 32 contact connectors (Omnetics) had been attached to the EIB. The tetrode tips were cut using fine sharp scissors (maximum length 3.5 mm and 3 mm for prelimbic cortex and HPC, respectively) and fixed to the inner tubes in the upper part. Tetrode tips were clean in distilled water and gold-plated (gold solution, Neuroalynx) using NanoZ software to lower their impedance to 100–200 kΩ and improve the signal-to-noise ratio. The tetrode tips were hidden at the same level as the bundle. The whole drive was covered with aluminum foil connected to the ground to reduce the electrostatic interference during the recordings. The bottom of the micro- drive was deepened in 70 % ethanol for 12 hours before brain implantation.

### Stereotaxic surgeries

#### Lentivirus injection

Lentiviral solutions were infused in the prelimbic cortex using stereotaxic surgery under biosafety level 2 conditions. The coordinates of the prelimbic cortex injection site were +3.2 mm AP, +/-0.8 mm ML from Bregma, and -2.5 mm DV from dura mater, according to The Rat Brain Atlas from Paxinos and Watson [45]. The procedure was carried out under isoflurane inhaled anesthesia. Unconsciousness was induced at 5 % isoflurane + 1 l/min O_2_ and maintained at 1.5-2 % isoflurane +1 l/min O_2_. A 0.8 mm diameter craniotomy was drilled above the target area in each hemisphere. The DV dura mater coordinate was measured before performing the durotomy.

A 30 G dental carpule connected to a 10 μl Hamilton and an infusion pump (Micro-pump, WPI) was slowly inserted into the brain target area (0.2 mm/min). A total volume of 2 μl of the lentiviral solution was infused at 200 nl/min. After 5 minutes of diffusion, the needle was removed, and the incision was sutured.

Temperature, oxygen saturation, and blood pressure were monitored during the whole surgical procedure. Some eye cream (Opthosam) was applied to protect the corneas during the intervention. At the start and end of the surgery, 2 ml of 0.9% NaCl physiological serum was administered subcutaneously. As analgesia, animals were administered 0.07 mg/ml carprofen in their water bottles two days before and three days after surgery. Immediately before surgery, 5 mg/kg carprofen was sc injected. In addition, a mix of 4 mg/kg lidocaine and 1 mg/kg bupivacaine in a 0.9% NaCl physiological serum was administered sc locally in the head.

After the viral injection, animals were housed individually in rat IVC cages for 14 days. Their weights and status were monitored daily for the correct recovery of animals. Then, rats were pair housed with their previous cagemate and moved to conventional housing.

#### Tetrode hyper-drive implantation

Twenty-one days after viral infusion, a second stereotaxic surgery took place for tetrode micro- drive implantation in 8 animals. The procedure was similar as described above. In this intervention additionally, a prophylactic 10 mg/kg sc injection of Baytril antibiotic was administered at the beginning of the surgery. Two craniotomies (2x1 mm and 1x1 mm for prelimbic cortex and HPC, respectively) were drilled above the target areas on the right hemisphere. The coordinates for the upper left corner of each craniotomy were: AP +4.5 mm and ML -0.5 (prelimbic cortex) and AP -3.8 mm and ML -2 mm (HPC) from Bregma [45]. A ground screw (M1x3) was placed on the left hemisphere in the cerebellum (AP -11 mm, ML +2 mm from Bregma). In addition, six M1x3 mm supporting screws were driven and bound to the skull using Super-bond C&B dental cement (Sun Medical, Japan). Carefully, the durotomies were performed, and the brain’s surface was exposed. Subsequently, the micro-drive was positioned on the brain’s surface, and attached to the skull and the screws by simplex rapid dental cement (Kemdent, UK). Then, tetrodes were slowly screw-driven into the prelimbic area in prelimbic cortex (3 mm DV from brain surface) and the cortical layers above the HPC (1.5 mm DV from brain surface). The dorsal hippocampal CA1 pyramidal layer was reached progressively in the subsequent days.

#### Object Space Task

The Object Space Task (OST) is a newly developed behavioral paradigm to study simple and semantic-like memories in rodents [13]. The task is based on the tendency of rodents to explore novel object-location in an open field across multiple trials. In these experiments, the OST took place as described previously [13] at least 21 days after the viral infusion when the effect of the RGS14_414_ protein is observed [40, 41]. Briefly, animals were handled for 5 consecutive days before and after surgery recovery. Then, rats were accommodated in the experimental room and habituated to the open field across 5 sessions (one per day). In the first session, animals explored the open field with their cagemate for 30 minutes. In the rest of the session, each individual freely explored the open field for 10 min. Two Duplo objects were included in the open field center in the last two sessions to facilitate a better exploration time in the subsequent task.

The OST consists of two phases: a training phase of 5 training trials in which animals are exposed to two identical objects (different across trials) for 5 minutes (45-55 min intertrial time); and a interference/test phase consisting of a single 10 min’ trial performed 24 h and 72 h after training. In the stable condition, both object locations were fixed during the training trials, and one object location was moved during the interference and tests sessions. In the overlapping condition, one object location was fixed, and the other one moved across training trials. In the interference and test sessions, the same object-location pattern from the last training trial was repeated.

The open field was a wooden square 75 x 75 x 60 cm. For the task, but not for the habituation, we placed 2D proximal cues on the open field walls and 3D distal cues above the open field. The cues were changed in each experimental session. In addition, the open field base colors changed across task sessions (white, blue, green, brown). The objects used vary in material (plastic, glass, wood, and metal), size, and colors. For electrophysiological recordings, not plastic objects were used to prevent static interference. All object bases were attached to 10 cm x 10 cm metal plates. Circular magnets were installed in the corners underneath the open field floor to prevent object movements during exploration. The open field and object surfaces were cleaned with 40% ethanol between trials to avoid odor biases.

Each trial was recorded using a camera above the open field. The object exploration time was manually scored *online* using the homemade software ’*Scorer32*’. The experimenter was blinded for treatment and experimental conditions at the moment of scoring. Object locations and experimental conditions were counterbalanced across treatments, individuals, and sessions.

For electrophysiological recordings, we run four cohorts of 2 animals each. Each cohort included one vehicle- and one RGS14_414_-treated animal, which performed the identical condition sequences with the same object-location patterns. Object locations and experimental conditions were counterbalanced across cohorts. Electrophysiological recordings took place during trials and rest periods (45 min before and 3 h after both training and test; and 45 min intertrial time during the training). Therefore, two brown wooden sleep boxes (40 x 75 x 60 cm height) with bedding material were placed next to the open field. Animals had been accustomed to the sleep box for at least 3 h in each open field habituation session.

Rats involved in the electrophysiological recordings also performed two experimental control conditions: homecage and random. The random condition was carried out as described previously [13], so there was a lack of repetitive object location patterns across different trials. In the homecage, the animal was recorded for 7 h and 10 min in the sleep box (a whole training session recording), and the experimenter kept the rat awake for the equivalent trial times.

#### In vivo electrophysiology recordings

In vivo freely moving extracellular recordings were executed during the OST and the resting periods. One session per experimental condition (homecage, stable, overlapping, and random) was carried out per animal. The local field potential (LFP) and single-unit activity detected by the 64 channels were amplified, filtered, and digitized through two 32 channels chip amplifier headstages (InstanTechnology) connected through the Intan cables and a commutator into the Open Ephys acquisition box. The signal was visualized using the open-source Open Ephys GUI (sample rate 30 kHz). In addition, the headstage contained an accelerometer to record the movement of the animals.

#### Sleep scoring

Different states (wakefulness, NonREM, REM, and Intermediate) were off-line manually scored using, ’TheStateEditor’ developed by Dr. Andres Grosmark at Dr. Gyorgy Buzsaki lab. One channel per brain area (Prelimbic cortex and hippocampus) was selected per animal. Using a 10 s sliding window, an experienced researcher created the hypnogram and 1 s epoch vector indicating brain states. The absences of movement in the accelerometer discriminate between wakefulness and sleep. During sleep periods, a dominant theta frequency in the dorsal hippocampus in the absence of spindles and delta waves indicated REM sleep. NonREM sleep was classified when slow oscillations were detected in the prelimbic cortex. The intermediate phase was defined as short transitional periods between NonREM and REM that show an increase in frequency in the prelimbic cortex and frequency similar to theta in the dorsal hippocampus. Microarousals were defined later on as periods of wakefulness <15 s within a sleep period.

#### Signal preprocessing

For the following analyses, first a single channel was selected per brain area. For prelimbic cortex, the channel with the largest slow oscillations was chosen. For hippocampus, the channel closest to the pyramidal layer, which displayed noticeable ripples was selected. Both channels were originally acquired at a sampling rate of 30 kHz and to avoid working which such a high rate, the channels were filtered with a 3rd order Butterworth lowpass filter at 500 Hz to avoid signal aliasing and then downsampled to 1 kHz.

#### Detection of spindles and delta waves

The downsampled prelimbic cortex channel (1 kHz) was loaded into the Matlab workspace and using a third-order Butterworth filter the signal was filtered to 9–20 Hz for detecting spindles and to 1–6 Hz for detecting delta waves. The NonREM bouts were then extracted from the filtered signal and concatenated. The functions FindSpindles and FindDeltaWaves from the Freely Moving Animal (FMA) toolbox http://fmatoolbox.sourceforge.net were modified and used to detect the start, peak and end of spindles and delta waves respectively. The optimal threshold was found for each animal by visually inspecting the detections and modifying the default parameters of the functions when needed. The results were saved as timestamps with respect to the concatenated NonREM signal in seconds. They were then used to find the timestamps with respect to the recorded signal. This process was repeated for pre and post trial sleep periods in study days pertaining to all animals in both treatment groups. The same method was applied for the detection of hippocampal delta waves.

#### Ripple detection

The downsampled channels (1 kHz) of the hippocampal pyramidal layer were loaded into the Matlab workspace and the NonREM bouts were extracted. Using a third-order Butterworth bandpass filter, the epochs of HPC signal were filtered to a frequency range of 100–300 Hz. A custom MATLAB function was used for detecting the start, peak and end of the ripples by thresholding voltage peaks which lasted a minimum duration of 30ms above the threshold. The start and end of the ripple were determined as half the value of the selected threshold. Values equivalent to 5 times the standard deviation of concatenated NonREM bouts were computed individually for presleep and all post trials in a study day. The average of these values was calculated to find a single detection threshold per study day. An offset of 5 microV was added to the threshold to reduce false positives. This was repeated for all study days pertaining to all animals in both treatment groups. To identify consecutive ripple events (multiplets), ripple peaks occurring within 200 ms of each other were grouped and classified as doublets, triplets, and so on, with their timestamps stored for further analysis.

#### Dimensionality reduction of ripple features and ripple clustering

Once ripples were detected, we extracted their waveforms (sampling rate = 1 kHz) using the start and end timestamps given by the detector. Instead of using a fixed temporal window, we computed ripple features using the intrinsic duration of each ripple waveform. Ripple features were computed as follows:

##### Average frequency

The Matlab function meanfreq() was used to calculate the mean frequency of ripples. It takes in two inputs, the signal, and the sample rate, and estimates the mean normalized frequency of the signal’s power spectrum. First, the power spectral density of the signal is computed using the periodogram() function and a default Kaiser window. Then the mean frequency is estimated as a weighted average, using the power spectral density values as weights of their corresponding frequencies.

##### Amplitude

To calculate the amplitude of the signal the envelope was computed by applying Hilbert transform. Then, the absolute value of this transform was taken to obtain real values. Finally, the maximum value of the envelope was calculated.

##### Area under the curve

The Matlab function trapz() was used to calculate the area under the curve of ripple waveforms absolute values. This function numerically integrated the absolute value of each ripple waveform over its duration in seconds.

##### Duration

The duration of each ripple was calculated in seconds using the start and end timestamps obtained from the ripple detector. As mentioned in the ripple detection section, the start and end of the ripple were determined as half the value of the amplitude detection threshold.

##### Peak-to-peak distance

The Matlab function peak2peak() was used to calculate the peak-to-peak distance. This function calculates the difference between the maximum and minimum amplitude values of the ripple waveform.

##### Power

The power of a ripple was computed as the squared Euclidean norm (i.e., vector magnitude) of a ripple waveform divided by its length in samples.

##### Entropy

The Matlab function entropy() was used to calculate the entropy of each ripple. Entropy is defined as -sum(p.*log2(p)), where p represents the normalized histogram counts.

##### Number of peaks

The Matlab function findpeaks() was used to find the local maxima (i.e., peaks), defined as data samples larger than its two neighboring samples without a minimum prominence. The total number of peaks was computed per ripple.

Each feature array was scaled using a z-score normalization. We performed principal component analysis (PCA) on the ripple features of aggregated ripples. For RGS14 ripples we used the PCA coefficient obtained from vehicle ripples to project the data into the same space and allow comparisons between treatments. We used the first three principal components which explained most of the variance. The resulting tridimensional point cloud was clustered using a Gaussian Mixture Model (GMM). To find the optimal number of clusters, Akaike information criterion (AIC) and Bayesian information criterion (BIC) were calculated from one cluster to ten clusters. AIC suggested four clusters as the best model while BIC five clusters. To verify further, the optimal number of clusters was calculated using k-means clustering and the Calinski-Harabasz criterion [46]. The result proved to be four. Therefore, the number of clusters was set to 4. Finally, the built-in MATLAB function fitgmdist() was used to fit a GMM to the data. The parameter ’CovarianceType’ was set to ‘’diagonal’.

#### Oscillation characteristics

The traces of each event detected (ripples, spindles, delta waves) were extracted using the start and end timestamps obtained from the detectors. The traces of the events were filtered in their corresponding detection frequency band. Characteristics such as the amplitude and mean frequency were calculated for these filtered events using built-in and custom MATLAB functions. Namely, the amplitude of the events was calculated by computing the envelope of the filtered trace using a Hilbert transform. The absolute value of the result was taken and its maximum was found. The mean frequency of the filtered traces was computed using the meanfreq function of MATLAB.

#### Detection of oscillation sequences

The sequences between ripples, spindles and delta waves were counted in various combinations to study cortico-hippocampal coupling during NonREM sleep as done by Maingret et al., 2016. The time difference between the peaks of these events was compared to a fixed duration to establish if there was a sequential relationship in the following combinations of oscillations: Delta-

Spindle (D-S), Delta-Ripple (D-R), Ripple-Delta (R-D), Ripple-Delta-Spindle (R-D-S), Delta-Ripple-Spindle (D-R-S). For D-S a sequence was considered when the duration between events was between 100– 1300ms, for D-R it was 50–400ms and for R-D it was 50–250ms. To find R-D-S sequences, the results of R-D and D-S were compared to find delta waves preceded by a ripple and followed by a spindle. To find D-R-S sequences, the results of D-R and R-S (events between 2-1000ms) were compared to find ripples preceded by a delta wave and followed by a spindle. To find the Delta-Spindle with Ripple sequences, the results of D-S and spindles co-occurring with ripples (see next subsection) were matched to find spindles preceded by a delta and co-occurring with a ripple. The results were saved as counts of each sequence for each post-trial.

#### Co-occurrence between ripples and spindles

The co-occurrence between ripples and spindles was computed by comparing the start and end timestamps of both events. To consider co-occurrence between a ripple and a spindle, either one of the following conditions had to be fulfilled. (1) A ripple had to start and end within the duration of the spindle. (2) One of the events had to start or end within the duration of the other. Given that more than one ripple can co-occur with the same spindle, we counted separately spindles co-occurring with spindles and spindles co-occurring with ripples.

#### Slow oscillation phase

The downsampled prelimbic cortex signal was filtered in the 0.5–4 Hz range using a third-order Butterworth bandpass filter and its Hilbert transform was computed to find the phase angle of slow oscillations in a range from 0° to 360°. The peaks of ripples and spindles were then used to find the corresponding slow oscillation phase. This same signal was later used to find the phase during spikes timestamps of cortical neurons.

#### Time-Frequency spectrograms

A two-second-long window centered on each ripple peak was extracted from the hippocampus and the prelimbic cortex channels respectively. All vehicle ripples across animals and conditions were combined for each ripple cluster and their amplitude was computed by finding the maximum of their envelope computed with a Hilbert transform. The median ripple amplitude was calculated and the corresponding two-second-long windows of the 2000 ripples which amplitude was the closest to the median amplitude were included in the following analysis. A notch filter at 50 Hz was applied to the ripple-centered windows using the ft_preprocessing function from the Matlab-based Fieldtrip toolbox Oostenveld et al., 2011. The Short-time Fourier transformation was calculated to detect the changes of spectral power in hippocampus and prelimbic cortex with respect to a time window of ±1 s around each ripple. This was computed using the ft_freqanalysis function from Fieldtrip with a 100ms Hanning window and time steps of 10ms, for a frequency range from 100 to 300 Hz with a 2 Hz step and from 0.5 to 20 Hz with a step of 0.5 Hz. The resulting spectrograms were averaged and displayed. To statistically compare spectrograms between ripple clusters, a nonparametric permutation test to correct for multiple comparisons with two-tailed pixel-based statistics was computed using 500 permutations and a p-value of 0.05 (Maris and Oostenveld, 2007).

#### Time-Frequency Granger causality

To determine the predictive power between brain regions during ripples, the time-frequency Spectral Granger Causality was computed for each directionality (Dhamala et al., 2008). A window with length of 2.2 s centered around each ripple peak was extracted for the simultaneous hippocampal and prelimbic cortex signals. The length of this window was chosen to at least capture one cycle of 0.5 Hz activity. A two-second-long time-frequency non-parametric Spectral Granger causality was computed by implementing a Short- time Fourier transform with a 500ms Hanning window with 10ms steps using the Fieldtrip functions ft_freqanalysis and ft_connectivityanalysis respectively. To determine statistical differences between Granger spectrograms, we created randomized trials by taking 400 random ripples per ripple cluster and computing their time-frequency Granger causality as described above. The result was stored, and the procedure was repeated 30 times to give a total of 30 randomized trials per ripple cluster. We then used the trials of the ripple clusters to determine significant statistical difference in each pixel of the time-frequency matrix by applying a nonparametric permutation test to correct for multiple comparisons with a two-tailed pixel-based correction, using 500 permutations and a p-value of 0.05.

#### Spike sorting

An important step of our analysis was identifying the cortical neurons and their spiking activity. The sleep and trial recordings of each tetrode of the prelimbic cortex were extracted and concatenated chronologically to allow tracking the individual neuronal activity across the whole study day. The tetrode recordings were preprocessed by applying a bandpass filter between 300 and 3000 Hz. For each tetrode, multiple spike sorting algorithms were run using the SpikeInterface python-based framework Buccino et al., 2020. The spike sorters used were Tridesclous, SpikingCircus, Klusta, HerdingSpikes, Ironclust, and MountainSort4. The default parameters included in SpikeInterface per spike sorter were used. Agreement between the spike sorters was computed and only putative neurons that were detected by at least two spike sorters were considered. When a tetrode didn’t have any consensus, the detections of the MountainSort4 spike sorter were used given that this spike sorter had better scores according to the SpikeForest Magland et al., 2020 metrics and our own examinations. After the putative neurons were detected, an experienced user curated manually the detections by visual inspection using the Phy interface and labeled them as either good, multiunit activity, or noise.

#### Spike waveform extraction

Sampled at 30 kHz, the prelimbic cortex tetrode channels were loaded and then filtered using a third-order Butterworth bandpass filter with a frequency range of 300–600 Hz. Neurons labeled as ‘good’ were used for further analysis. A preliminary waveform per neuron was defined with an 82- sample window with 40 samples before and 41 samples after the spike timestamp (ST). For each neuron, STs were randomly permuted and a total of 2000 were selected. The permutation was done in order to avoid bias when selecting waveforms. These 2000 waveforms were averaged to obtain the mean spike waveform per neuron. Since tetrodes were used to record neuronal activity during the task and sleep, there were 4 average waveforms to choose from. The one with the highest peak amplitude was chosen. For each neuron, all STs were stored. Further, each neuron was assigned a unique ID. This process was performed for all neurons of all rats.

#### Neuron classification

The average neuron waveforms calculated from rats 1, 2, 6, and 9 were categorized as ‘Vehicle’ and stored in a 139 (t1) ×82 matrix, with 139 being the total number of putative neurons and 82 being the number of samples as mentioned before. Similarly, ‘RGS14’ data from rats 3, 4, 7, 8 was stored as a 353 (t2) ×82 matrix. Variables t1 and t2 were the total counts of neurons for the respective treatments. The bz_CellClassification.m script from Buzsaki lab’s GitHub was used to compute the trough-to-peak delay time and the spike width for the spike waveform of each putative neuron. The function ClusterPointsBoundaryOutBW.m of the same repository was modified to incorporate visualization of interneuron and pyramidal data for both treatments. Neurons were then visualized in a 2D plane with x and y axes as trough-to-peak delay and spike width respectively. The set of coordinates of all putative neurons were fed into a Gaussian Mixture Model (GMM) with two components in order to find the centroids of the clusters of pyramidal cells and interneurons. The cluster which contained spikes with high values of spike width and trough-to-peak delay was labeled as the pyramidal neurons cluster, while the remaining one was labeled as the interneurons cluster. For both clusters, a threshold of mean +/- 2 SD with respect to their centroids was used to filter out the outliers. Next, the firing rates of the remaining neurons were calculated as is described in the following paragraph and those with extreme firing rate values were reinspected in the Phy interface by another experimenter to potentially discard remaining false positives. After this procedure there were a total of 101 pyramidal neurons for the RGS14 treatment and 57 for Vehicle. The total number of interneurons were 18 for RGS14 and 7 for Vehicle. In the following analyses, only pyramidal neurons were used given the low number of interneurons detected.

#### Cortical activity during ripples

Using the spike timestamps during NonREM sleep, the cortical pyramidal neuron response to hippocampal ripples was computed in a 2 s window defined around the peak of each ripple. For each ripple the activity of all pyramidal neurons detected during that day was extracted. The spikes timestamps in the 2 s window were normalized to vary from –1 to 1 s, where 0 was the ripple peak. After concatenating the normalized timestamps across all ripple-centered windows per neuron, they were binned in 10ms bins and the number of spikes was determined for each bin. Hence, for each neuron, a [1x200] column vector was obtained. This vector was then z-normalized. The final z-scored vector was found by averaging the z-scored vector over all neurons. The data was smoothed twice using the MATLAB smooth function. To visualize the activity of prelimbic cortex pyramidal neurons around the ripple, the firing activity of a neuron was determined from a 50ms window around the peak of the ripple (–20 to +30ms) by quantifying the number of spikes. After compiling the firing activity around the ripple peak for each neuron the values were sorted in an ascending order and was visualized using the MATLAB imagesc function.

---

**SFig. 1.**
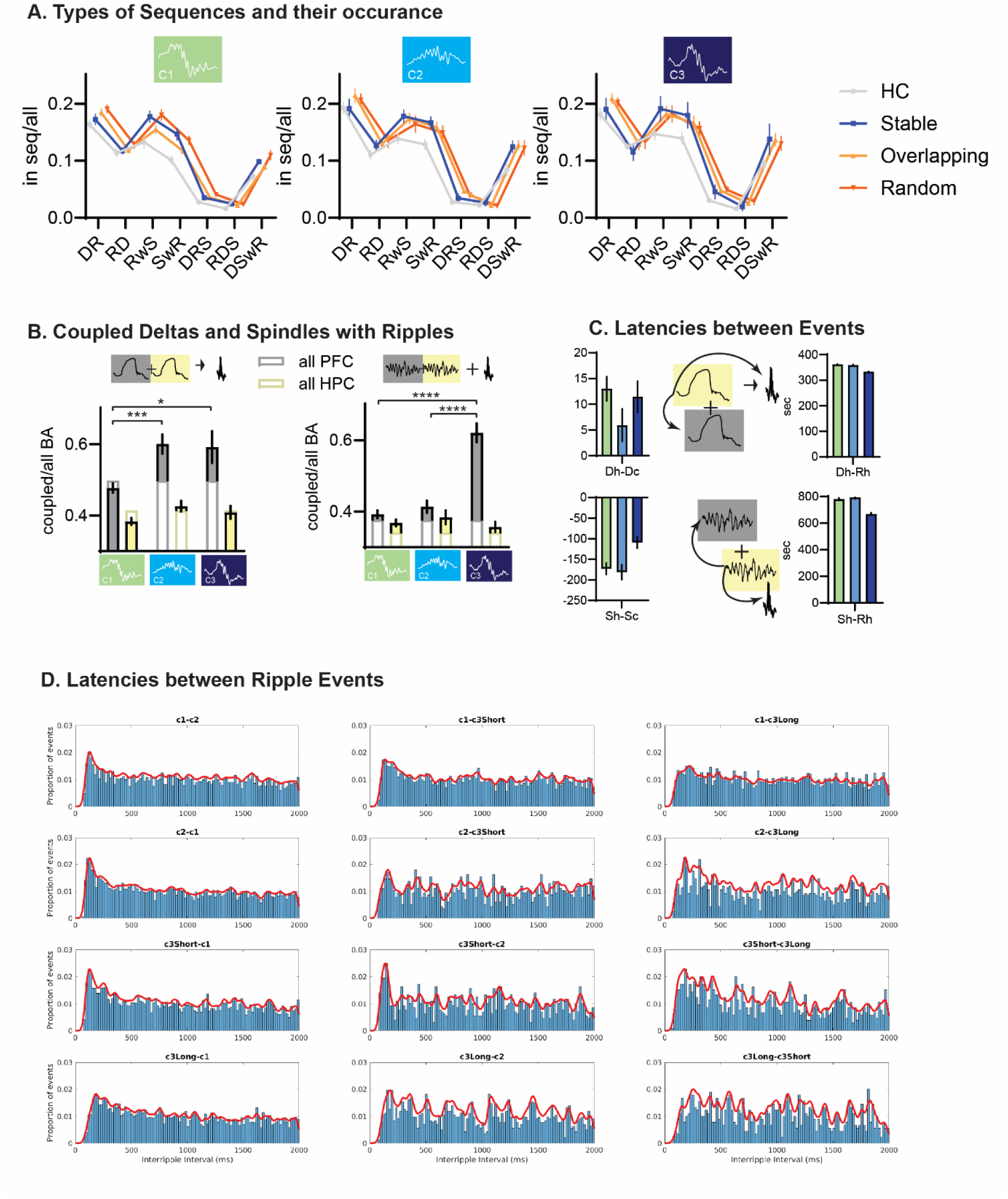
Sequences and Couplings: A. For each cluster type the different sequences and their occurrence rates split for conditions. All types of clusters increased DR coupling in the complex memory conditions and ripple during spindle (RwS), spindle with at least one ripple (SwR), and the same with a preceding delta (DSwR) all increased for simple and complex learning. Findings already reported in [47]. B. Fraction of delta and spindle (left, right respectively) that were coupled to their hippocampal or cortical counterparts considering DR and SwR events. For DR C2 and C3 showed higher cortical to hippocampal coupling than baseline (simple D-D and S-S events) and for SwR only C3 showed this increased coupling of cortical events. C. Latencies of these coupled events. D. Latencies of ripple type succession. C3short-C2 latencies had a more distinct peak at the expected doublet/multiplet interval

**SFig. 2.**
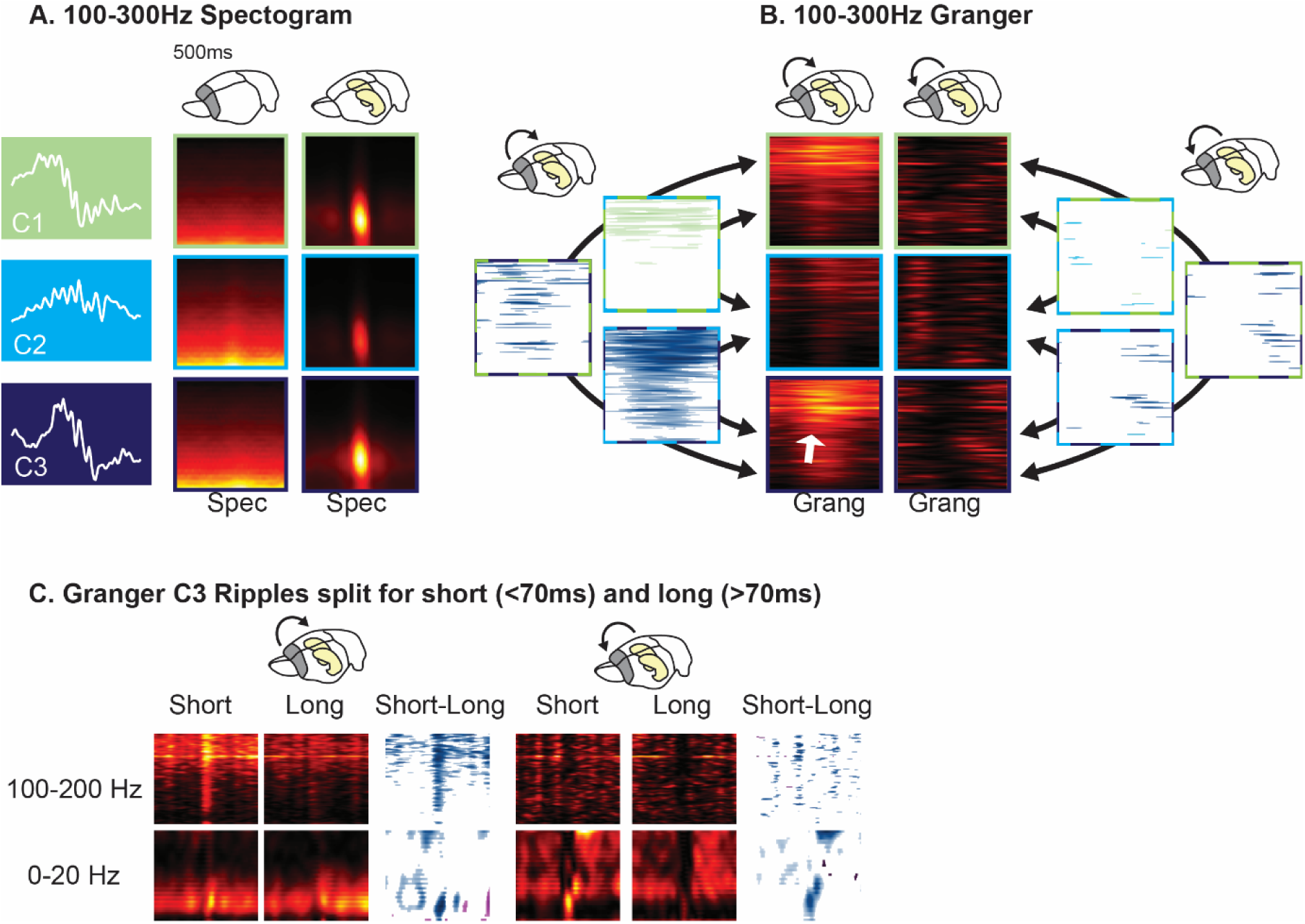
Spectrograms and Granger Causality Analysis. A and B for 100-300 Hz, C Granger split for short and long C3.

**SFig. 3.**
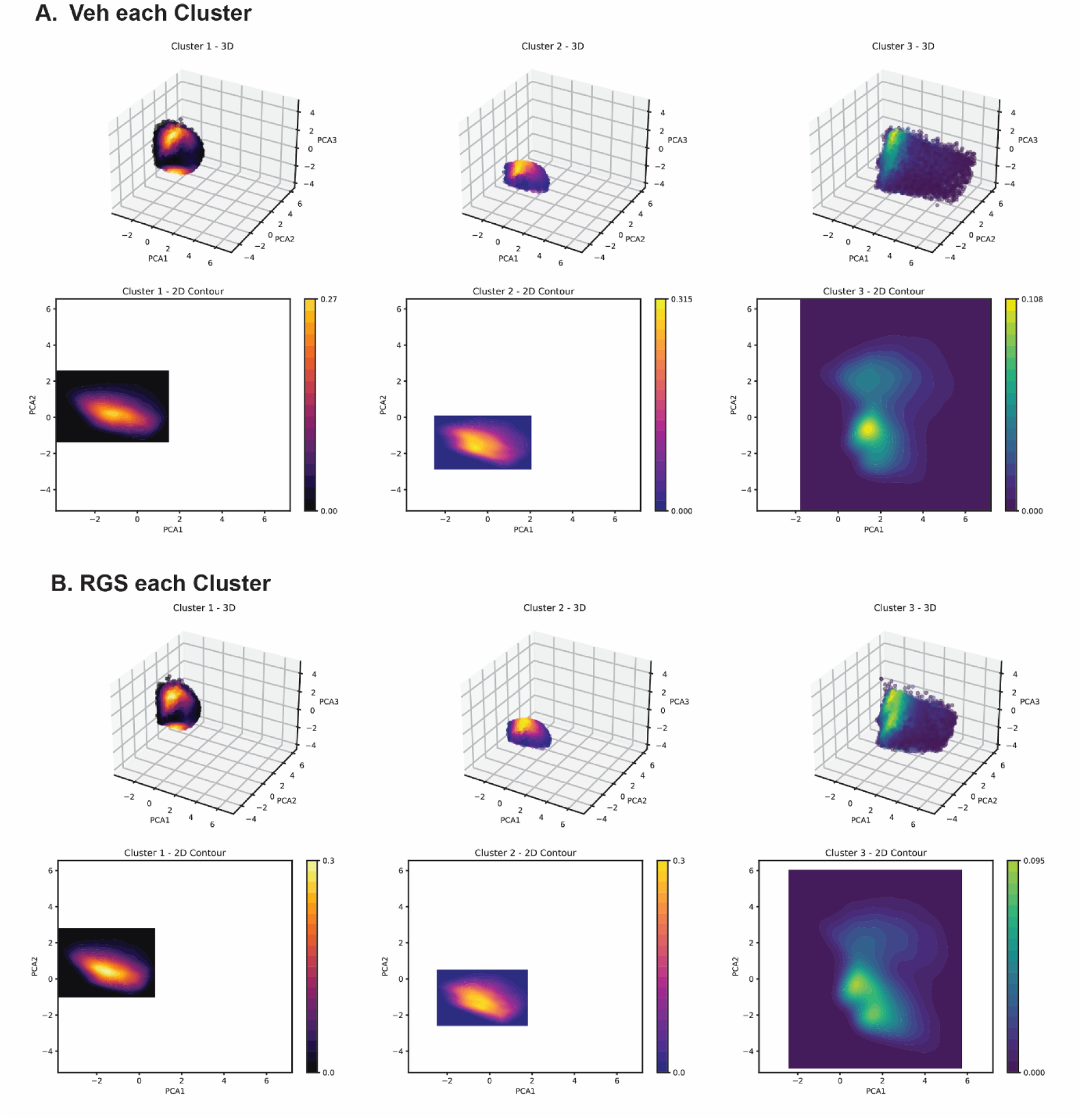
Ripple-PCA in Veh and RGS14 each cluster.

**SFig. 4.**
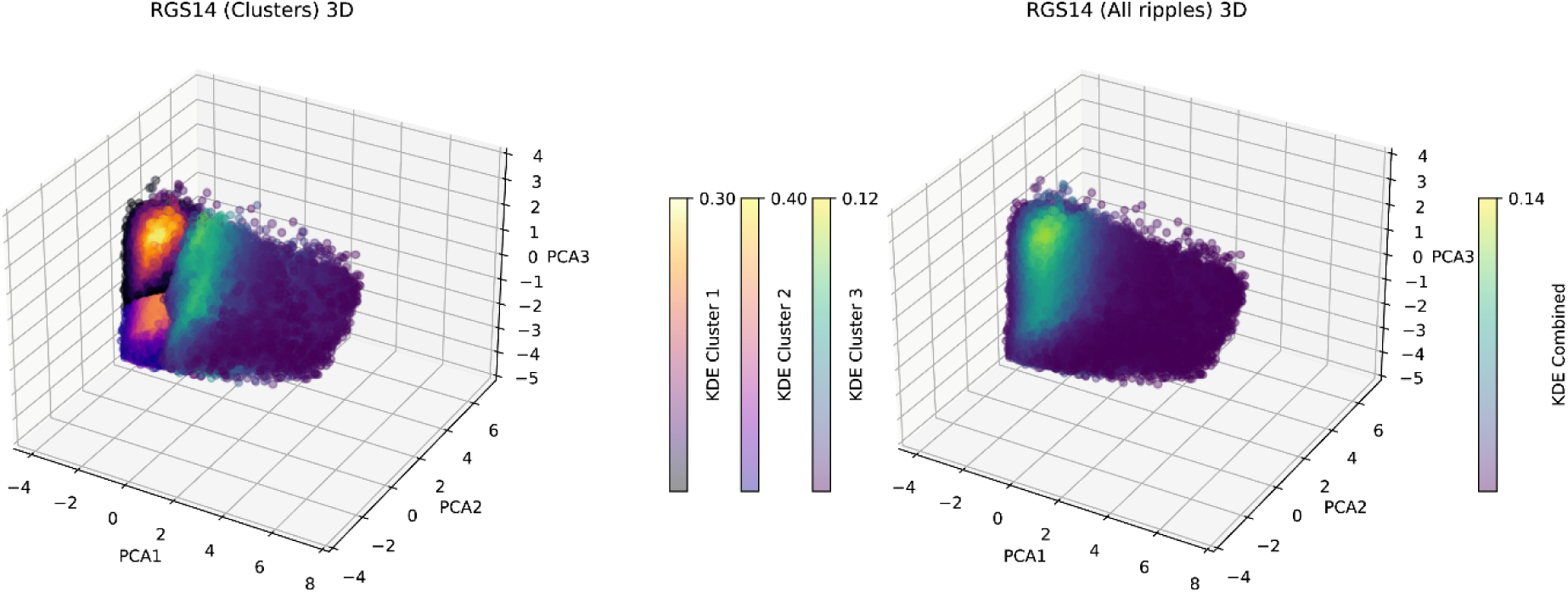
Ripple-PCA in RGS14 with and without clusters.

SVideo 1: Rotating 3D view of ripples in PCA space with and without clusters.

SVideo 2: Rotating 3D view of ripples in PCA space split for clusters

SVideo 3: Rotating 3D view of ripples in PCA space split for clusters in RGS14 treated animals.

## References

1. Girardeau, G., et al., Selective suppression of hippocampal ripples impairs spatial memory. Nat Neurosci, 2009. 12(10): p. 1222–3.

2. Ego-Stengel, V. and M.A. Wilson, Disruption of ripple-associated hippocampal activity during rest impairs spatial learning in the rat. Hippocampus, 2010. 20(1): p. 1–10.

3. Aleman-Zapata, A., R.G.M. Morris, and L. Genzel, Sleep deprivation and hippocampal ripple disruption after one-session learning eliminate memory expression the next day. Proceedings of the National Academy of Sciences, 2022. 119(44): p. e2123424119.

4. Navarro-Lobato, I., et al., Increased cortical plasticity leads to memory interference and enhanced hippocampal-cortical interactions. Elife, 2023. 12:e84911.

5. Bendor, D. and M.A. Wilson, Biasing the content of hippocampal replay during sleep. Nat Neurosci, 2012. 15(10): p. 1439–1444.

6. McNamara, C.G., et al., Dopaminergic neurons promote hippocampal reactivation and spatial memory persistence. Nat Neurosci, 2014. 17(12): p. 1658–60.

7. Zutshi, I. and G. Buzsáki, Hippocampal sharp-wave ripples and their spike assembly content are regulated by the medial entorhinal cortex. Curr Biol, 2023. 33(17): p. 3648–3659.e4.

8. Oliva, A., et al., Hippocampal CA2 sharp-wave ripples reactivate and promote social memory. Nature, 2020. 587(7833): p. 264-269.

9. Stark, E., et al., Pyramidal cell-interneuron interactions underlie hippocampal ripple oscillations. Neuron, 2014. 83(2): p. 467–80.

10. Sebastian, E.R., et al., Topological analysis of sharp-wave ripple waveforms reveals input mechanisms behind feature variations. Nature Neuroscience, 2023. 26(12): p. 2171–2181.

11. Ramirez-Villegas, J.F., N.K. Logothetis, and M. Besserve, Diversity of sharp-wave-ripple LFP signatures reveals differentiated brain-wide dynamical events. Proc Natl Acad Sci U S A, 2015. 112(46): p. E6379–87.

12. Aleman-Zapata, A., et al., Differential Contributions of CA3 and Entorhinal Cortex Inputs to Ripple Patterns in the Hippocampus Under Cannabidiol. bioRxiv, 2024: p. 2024.08.06.606645.

13. Genzel, L., et al., The object space task shows cumulative memory expression in both mice and rats. PLoS Biol, 2019. 17(6).

14. Molle, M., et al., Grouping of spindle activity during slow oscillations in human non-rapid eye movement sleep. J Neurosci, 2002. 22(24): p. 10941–7.

15. Peyrache, A., et al., Replay of rule-learning related neural patterns in the prefrontal cortex during sleep. Nat Neurosci, 2009. 12(7): p. 919–926.

16. Maingret, N., et al., Hippocampo-cortical coupling mediates memory consolidation during sleep. Nature Neuroscience, 2016. 19(7): p. 959–64.

17. Sirota, A., et al., Communication between neocortex and hippocampus during sleep in rodents. Proceedings of the National Academy of Sciences, 2003. 100(4): p. 2065–2069.

18. Diekelmann, S. and J. Born, The memory function of sleep. Nat Rev Neurosci, 2010. 11(2): p. 114–126.

19. Samanta, A., et al., CBD lengthens sleep but shortens ripples and leads to intact simple but worse cumulative memory. iScience, 2023. 26(11): p. 108327.

20. Caragea, V.-M. and D. Manahan-Vaughan, Bidirectional Regulation of Hippocampal Synaptic Plasticity and Modulation of Cumulative Spatial Memory by Dopamine D2-Like Receptors. Frontiers in Behavioral Neuroscience, 2022. 15.

21. Rasch, B. and J. Born, About sleep’s role in memory. Psychol Rev, 2013. 93: p. 681–766.

22. Navarro-Lobato, I., et al., 14-3-3ζ is crucial for the conversion of labile short-term object recognition memory into stable long-term memory. J Neurosci Res, 2021. 99(9): p. 2305–2317.

23. Masmudi-Martín, M., et al., RGS14(414) treatment induces memory enhancement and rescues episodic memory deficits. Faseb j, 2019. 33(11): p. 11804–11820.

24. El-Kanbi, K., et al., Distinction between slow waves and delta waves sheds light to sleep homeostasis and their association to hippocampal sharp waves ripples. bioRxiv, 2022: p. 2022.12.27.522034.

25. Schreiner, T., et al., The human thalamus orchestrates neocortical oscillations during NREM sleep. Nature Communications, 2022. 13(1): p. 5231.

26. Sullivan, D., et al., Relationships between hippocampal sharp waves, ripples, and fast gamma oscillation: influence of dentate and entorhinal cortical activity. J Neurosci, 2011. 31(23): p. 8605–16.

27. Fernández-Ruiz, A., et al., Long-duration hippocampal sharp wave ripples improve memory. Science, 2019. 364(6445): p. 1082-1086.

28. Nitzan, N., et al., Brain-wide interactions during hippocampal sharp wave ripples. Proc Natl Acad Sci U S A, 2022. 119(20): p. e2200931119.

29. Pochinok, I., et al., A developmental increase of inhibition promotes the emergence of hippocampal ripples. bioRxiv, 2023.

30. Schreiner, T., M. Lehmann, and B. Rasch, Auditory feedback blocks memory benefits of cueing during sleep. Nat Commun, 2015. 6: p. 8729.

31. Hall, A.F. and D.V. Wang, The two tales of hippocampal sharp-wave ripple content: The rigid and the plastic. Prog Neurobiol, 2023. 221: p. 102396.

32. Grosmark, A.D. and G. Buzsaki, Diversity in neural firing dynamics supports both rigid and learned hippocampal sequences. Science, 2016. 351(6280): p. 1440-3.

33. Girardeau, G., A. Cei, and M. Zugaro, Learning-induced plasticity regulates hippocampal sharp wave-ripple drive. J Neurosci, 2014. 34(15): p. 5176–83.

34. Domanski, A.P.F., et al., Distinct hippocampal-prefrontal neural assemblies coordinate memory encoding, maintenance, and recall. Curr Biol, 2023. 33(7): p. 1220–1236.e4.

35. Isomura, Y., et al., Integration and Segregation of Activity in Entorhinal-Hippocampal Subregions by Neocortical Slow Oscillations. Neuron, 2006. 52(5): p. 871–882.

36. Staresina, B., et al., Hierachical nesting of slow oscillations, spindles and ripples in the human hippocampus during sleep. Nat Neurosci, 2015. 18(11): p. 1679–86.

37. Genzel, L., et al., Light sleep versus slow wave sleep in memory consolidation: a question of global versus local processes? Trends in Neurosciences, 2014. 37(1): p. 10–19.

38. Bozic, I., et al., Coupling between the prelimbic cortex, nucleus reuniens, and hippocampus during NREM sleep remains stable under cognitive and homeostatic demands. Eur J Neurosci, 2023. 57(1): p. 106–128.

39. Ferraris, M., et al., The nucleus reuniens, a thalamic relay for cortico-hippocampal interaction in recent and remote memory consolidation. Neuroscience & Biobehavioral Reviews, 2021. 125: p. 339–354.

40. Lopez-Aranda, M.F., et al., Role of layer 6 of V2 visual cortex in object-recognition memory. Science, 2009. 325(5936): p. 87-9.

41. Masmudi-Martin, M., et al., RGS14414 treatment induces memory enhancement and rescues episodic memory deficits. FASEB J, 2019. 33(11): p. 11804–11820.

42. Brunetti, P.M., et al., Design and fabrication of ultralight weight, adjustable multi-electrode probes for electrophysiological recordings in mice. J Vis Exp, 2014(91): p. e51675.

43. Kloosterman, F., et al., Micro-drive array for chronic in vivo recording: drive fabrication. J Vis Exp, 2009(26).

44. Nguyen, D.P., et al., Micro-drive array for chronic in vivo recording: tetrode assembly. J Vis Exp, 2009(26).

45. Paxinos, G.W., Charles, The Rat Brain Atlas in Stereotaxic Coordinates. 7th Edition ed. 2013: Academic Press. 472.

46. Caliński, T. and J. Harabasz, A dendrite method for cluster analysis. Communications in Statistics, 1974. 3(1): p. 1–27.

47. Navarro Lobato, I., et al., Increased cortical plasticity leads to memory interference and enhanced hippocampal-cortical interactions. eLife, 2023. 12: p. e84911.

